# HER3 mediates metabolic reprogramming in metastatic colorectal and pancreatic cancer

**DOI:** 10.64898/2026.10.30.763164

**Authors:** Moeez Rathore, Chao Wei, Kimberly Curry, Mehrdad Zarei, Hallie Graor, Jordan Winter, Rui Wang

**Affiliations:** Department of Surgery, Case Western Reserve University, 10900 Euclid Ave., Cleveland, OH 44106; Case Comprehensive Cancer Center, Case Western Reserve University, 10900 Euclid Ave., Cleveland, OH 44106; Department of Surgery, Division of Surgical Oncology, University Hospitals Cleveland Medical Center, 11100 Euclid Avenue, Cleveland, OH 44106, USA

**Keywords:** CRC, PDAC, glycolysis, HER3, PFK2, OXPHOS

## Abstract

Liver metastases are a major cause of mortality in colorectal cancer (CRC) and pancreatic ductal adenocarcinoma (PDAC), yet the oncogenic signals that shape tumor metabolism in the hepatic niche remain poorly understood. Here, we show that HER3 activation drives a liver metastasis-specific glycolytic program in both cancer types. Genetic or pharmacological HER3 inhibition suppresses cancer glycolysis while increasing dependence on oxidative phosphorylation (OXPHOS), rendering liver metastases vulnerable to OXPHOS inhibition. Combining the HER3 inhibitor sapitinib with distinct OXPHOS inhibitors led to *in vitro* synergy and durable tumor-free survival in orthotopic CRC and PDAC liver metastases models, with further validation in patient-derived xenografts. Mechanistically, HER3 promotes glycolysis by activating PFK2 and reduces OXPHOS by activating Drp1-mediated mitochondrial remodeling; both pathways were activated in liver metastases compared with matched primary tumors. Together, our findings identify the HER3–AKT/RSK–PFK2 axis as a driver of glycolysis and metastatic growth and reveal an OXPHOS dependency induced by HER3 inhibition that may be exploited through combined HER3 and OXPHOS targeting in metastatic CRC and PDAC.

## Introduction

The liver is the most frequent site of distant metastasis in colorectal cancer (CRC) and pancreatic ductal adenocarcinoma (PDAC), and liver metastasis is a major determinant of mortality in both diseases [1, 2]. Despite advances in systemic therapy, patients with liver-metastatic CRC (mCRC) and PDAC (mPDAC) continue to have poor outcomes, with median overall survival measured in months [3, 4]. Progress toward more effective therapies is limited, in part, by an incomplete understanding of how the hepatic microenvironment reshapes tumor cell biology to support metastatic growth.

The hepatic microenvironment differs metabolically from primary tumor sites and contains abundant glucose, fatty acids, and amino acids that may differentially support metastatic outgrowth. Emerging evidence indicates that cancer cells undergo substantial metabolic reprogramming upon colonization of the liver [5, 6]. However, the oncogenic signals that enable this adaptation in CRC and PDAC liver metastases remain poorly defined. Identifying these signals may reveal therapeutic vulnerabilities specific to the metastatic site.

HER3 (ERBB3), a member of the ERBB receptor tyrosine kinase family, has emerged as an important mediator of metastatic progression in several gastrointestinal cancers [7, 8]. HER3 ligands, including neuregulin-1 (NRG1) and leucine-rich alpha-2-glycoprotein 1 (LRG1), are abundant in the hepatic microenvironment, suggesting that liver colonization may selectively activate HER3 signaling in disseminated tumor cells [9]. Hepatocyte-derived LRG1 has also been shown to prime the liver for metastasis and impair immunotherapy [10]. Consistent with a role for this pathway, HER3 expression is increased in CRC liver metastases relative to matched primary tumors [11]. However, whether HER3 mediates metabolic reprogramming in liver metastases, and whether this creates therapeutically exploitable vulnerabilities, remains unknown.

Here, using CRC and PDAC liver metastasis models, we show that HER3 drives a hepatic niche-specific metabolic program characterized by increased glycolysis and reduced oxidative phosphorylation (OXPHOS), thereby supporting metastatic outgrowth. We identify a HER3– RSK/AKT–PFK2 signaling axis that promotes glycolysis and a parallel HER3–DRP1 S616 pathway associated with reduced OXPHOS. HER3 inhibition exposes adependence on OXPHOS in liver metastatic cells, and combined targeting of HER3 and OXPHOS produces durable tumor-free survival in orthotopic mCRC and mPDAC models, with supporting results in patient-derived xenografts. These findings provide a mechanistic framework for HER3-mediated metabolic adaptation in the liver and a rationale for combination therapy in metastatic gastrointestinal cancers.

## Results

### HER3 ablation suppresses glycolysis in liver metastases and exposes OXPHOS dependency

To determine whether HER3 activation changes during liver metastasis, we assessed HER3 by immunohistochemistry (IHC) in paired primary tumors and corresponding liver metastases. Compared to primary tumors, HER3 phosphorylation (Y1289) was markedly elevated in matched liver metastases from the same patient, consistent with selective induction or enrichment within the hepatic microenvironment **(Fig. 1A)**. We next used orthotopic murine pancreatic cancer models to establish primary tumors and liver tumors by pancreatic or intrahepatic injecting KPC cells, respectively. Untargetedmetabolomic profiling showedincreased glycolytic intermediates in PDAC liver metastases relative to primary tumors **(Fig. 1B)**. To examine the role of *HER3* genetically, we used an orthotopic KPC liver tumor model with *HER3* knockout (HER3-KO). Targeted liquid chromatography–mass spectrometry (LC-MS) profiling revealed a coordinated reduction in central carbon metabolic intermediates in HER3-KO tumors compared with HER3-WT tumors (**Fig. 1C, D**). To systematically define the global metabolic consequences of HER3 depletion, we performed metabolite set enrichment analysis (MSEA) comparing HER3-WT and HER3-KO cells. Unbiased pathway profiling of the top 25 enriched metabolite sets revealed a coordinated metabolic shift spanning redox defense, nitrogen and amino acid assimilation, and core bioenergetic pathways **(Fig. 1E)**. Together, these pathway shifts indicate that loss of HER3 suppresses glycolytic flux while driving a compensatory upregulation of mitochondrial metabolism.

**Figure 1.**
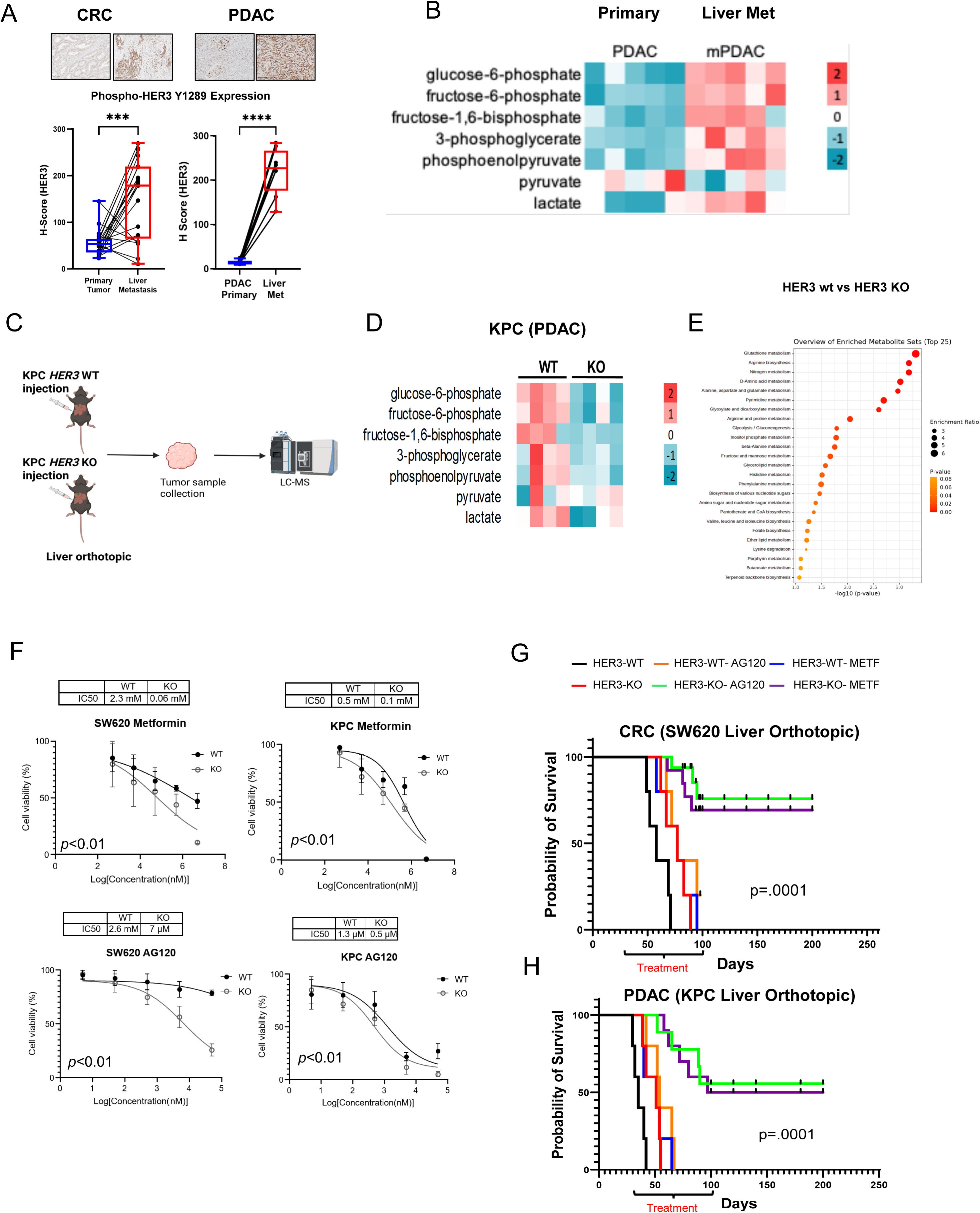
HER3 blockade creates a metabolic dependency that sensitizes metastatic CRC and PDAC to OXPHOS inhibition. (A) Immunohistochemical (IHC) staining and quantification of HER3 phosphorylation (S1289) in human CRC and PDAC matched primary tumors (blue) and liver metastases (red). Each dot represents one biologically independent human sample. (B) Heatmap shows metabolite levels in KPC murine PDAC orthotopic primary tumors and liver metastases. (C) Schematics of developing and analyzing KPC murine orthotopic PDAC liver metastases (D and E) Heatmap and pathway analysis of metabolite changes between wild-type (WT) or HER3-knckout (KO) KPC tumors with (F) Relative growth and IC50 of WT and HER3 KO cells responses to metformin or the IDH1 inhibitor AG120. (G and H) Kaplan–Meier overall survival of mice bearing WT or HER3 KO orthotopic liver metastases and treated with either metformin (METF) or AG120. Data in presented as mean ± SEM. \*\*\**p<0.0001,* \*\*\*\**p<0.00001,* by t-test. *p* values in F determined by t-test, *p* values in G determined by log-rank test.

To test whether HER3 loss increases dependence on OXPHOS *in vitro*, we treated PDAC and CRC cell lines with the OXPHOS-targeting agents metformin [12], AG120 [13], and two additional clinically relevant OXPHOS inhibitors IACS-010759 [14], lixumistat hydrochloride [15]. Dose–response viability assays showed substantially lower IC50 values in HER3 KO cells than in WT cells, consistent with increased sensitivity to these agents following HER3 loss **(Fig. 1F, Supp Fig. 1)**. Thus, HER3 ablation reveals a metabolic vulnerability to OXPHOS inhibition in both cancer types.

We next tested this vulnerability *in vivo* using orthotopic liver metastasis models generated with HER3 WT or KO cells and treated with AG120 or metformin. Kaplan–Meier analysis showed that HER3 KO or inhibitor alone modestly prolonged survival. In contrast, combining HER3 KO with either inhibitor significantly extended survival in the mPDAC (KPC) and mCRC (SW620) models **(Fig. 1G, H).** Approximately 50% of animals achieved durable tumor-free survival after treatment discontinuation.

### Combined HER3 and OXPHOS inhibition achieves tumor-free survival in liver metastasis models

Consistent with our *in vitro* metabolite enrichment analysis showing that genetic HER3 loss suppresses glycolysis and shifts cellular bioenergetics toward mitochondrial metabolism, in KPC mice treated with the HER3 inhibitor sapitinib **(Fig. 2A)**, targeted LC-MS metabolomics revealed that untreated liver metastases exhibit elevated glycolytic intermediates and lactate relative to primary tumors, whereas pharmacologic HER3 inhibition attenuates glycolytic intermediate and lactate accumulation, most notably in liver metastases **(Fig. 2B)**. Unbiased metabolite set enrichment analysis comparing CTRL versus HER3 inhibitors further demonstrated significant remodeling of core mitochondrial and amino acid-linked pathways (**Supp Fig. 2A)** — including pyruvate metabolism, the citrate cycle (TCA cycle), glyoxylate and dicarboxylate metabolism, and histidine metabolism—collectively showing that HER3 blockade suppresses glycolysis while driving a metabolic shift toward mitochondrial and TCA-coupled pathways.

**Figure 2.**
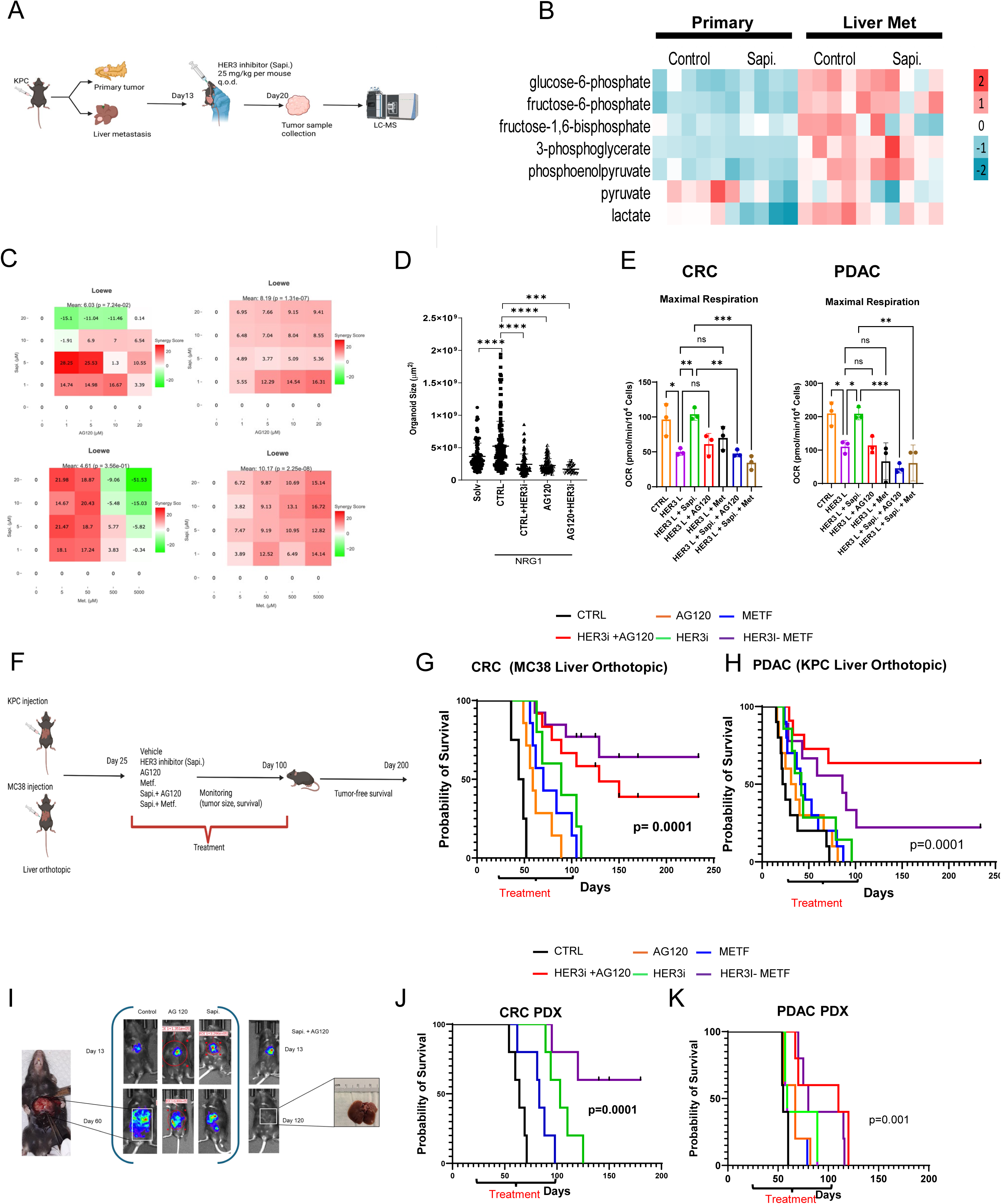
Pharmacological Inhibitions of HER3 and OXPHOS lead to in vitro synergy and durable complete responses in animal models. (A and B) Schematics of developing KPC murine orthotopic PDAC liver metastases and drug treatments, with resulting changes in metabolites shown by heatmap. (C) Loewe additivity model to determine *In vitro* synergy of combining HER3 inhibitors sapitinib (Sapi) and OXPHOS inhibitors, either IDH1 inhibitors AG120 or metformin (Met.). (D) Sizes of CRC organoids cultured with indicated treatments. Each dot represents one organoid, pooled from three independent experiments using the same organoid line. (E) Oxygen consumption rate (OCR) determined by Seahorse FX analysis. (F—H) Schematics of developing orthotopic liver metastases and drug treatments, with resulting mouse survival determined by Kaplan–Meier. (I) Representative pictures of mice with tumor-bearing liver and *in vivo* bioluminescence. (J and K) Kaplan–Meier overall survival of mice with PDXs. Data in D and E presented as mean ± SEM. \*\**p<0.001,* \*\*\**p<0.0001,* \*\*\*\**p<0.00001,* ns, not significant by t-test. *p* values in G, H, J and K are determined by log-rank test.

Given HER3’s effects on glycolysis and OXPHOS, we next assessed the therapeutic potential of combined HER3 and OXPHOS inhibition. A HER3 inhibitor (HER3i) combined with AG120 or metformin showed strong synergy in CRC and PDAC cell lines **(Fig. 2C, Supp Fig. 2B).** These effects were further confirmed in additional cell lines and patient-derived CRC organoids **(Fig. 2D, Supp Fig. 2C).** Seahorse oxygen consumption rate (OCR) analysis showed that combining HER3i with AG120 or metformin suppressed mitochondrial respiration more effectively than either agent alone **(Fig. 2E, Supp Fig. 2D).**

To evaluate efficacy *in vivo,* we treated orthotopic liver metastasis models (MC38 for CRC and KPC for PDAC) with sapitinib, AG120, metformin, or the corresponding combinations **(Fig. 2F)**. Combined HER3 and OXPHOS inhibition significantly prolonged survival compared with either agent alone **(Fig. 2G, H).** Bioluminescence imaging demonstrated tumor regression with combination treatment, accompanied by complete tumor-free survival after treatment discontinuation **(Fig. 2I).** We further validated these findings in patient-derived xenograft (PDX) models of CRC and PDAC, in which the combinations significantly extended survival and reduced tumor volume compared with controls **(Fig. 2J, K)**. Together, these data support the therapeutic potential of combining HER3 and OXPHOS targeting in CRC and PDAC liver metastases.

### HER3 promotes glycolysis through PFK2

To investigate how HER3 promotes glycolysis, we first examined the effects of the HER3 ligands NRG1 and LRG1. Stimulation with either ligand increased lactate production in CRC and PDAC cell lines **(Fig. 3A).** Functionally validating our metabolomic profiling, real-time longitudinal measurements of extracellular metabolites in KPC cells over 60 hours revealed that genetic deletion of HER3 significantly impairs both glucose consumption and lactate secretion, confirming a direct requirement for HER3 in sustaining glycolysis **(Fig. 3B).**

**Figure 3.**
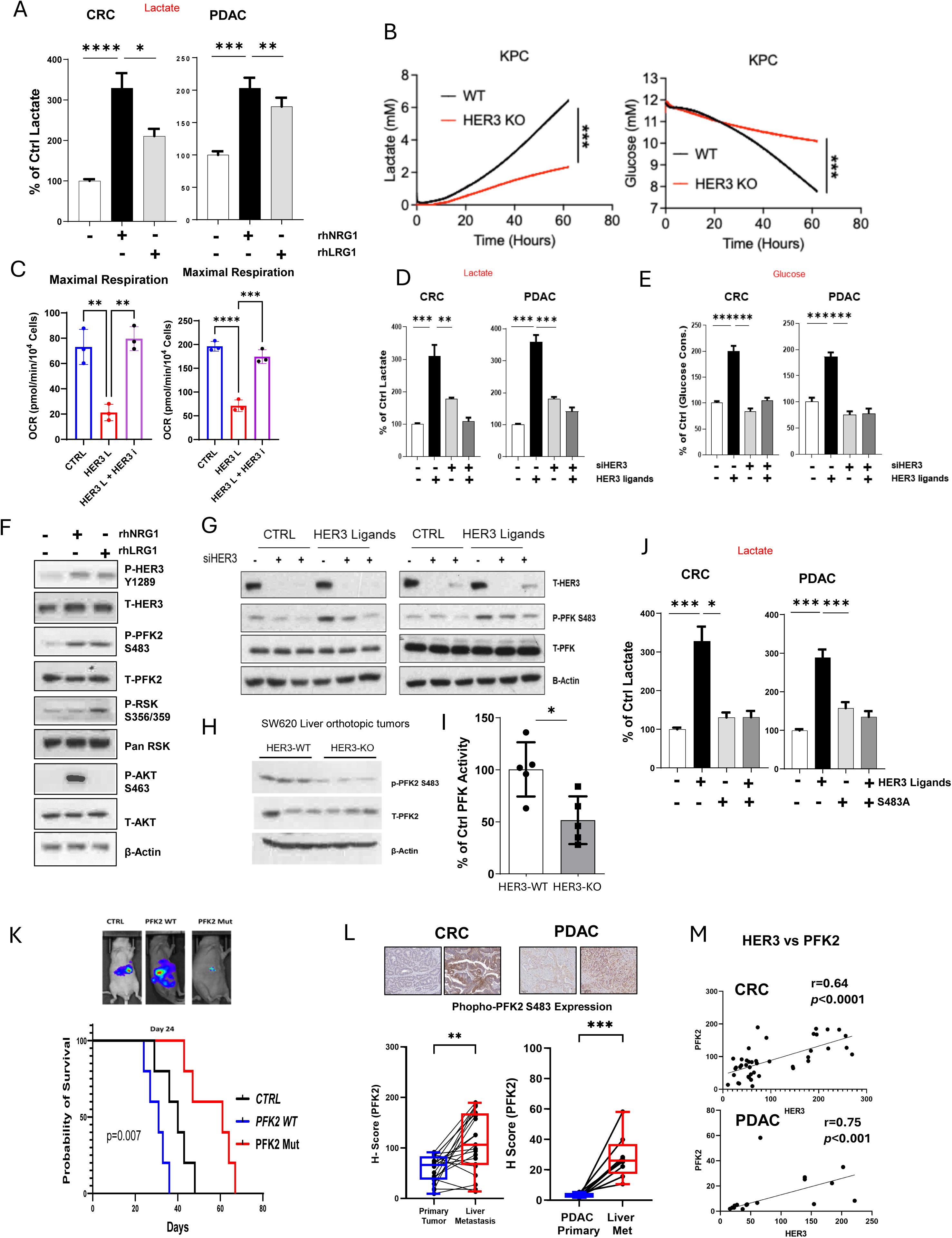
PFK2 signaling mediates HER3-induced glycolysis in cancer cells. (A and B) Lactate secretion determined by ELISA. (C) Oxygen consumption rate (OCR) determined by Seahorse FX analysis. (D) Lactate secretion determined by ELISA. (E) Glucose consumption determined by ELISA. (F and G) Western blotting showing levels of indicated proteins in SW620 human CRC cells with desired treatments. (H and I) Western blotting and ELISA assays showing levels of indicated proteins and PKA kinase activity in orthotopic CRC liver metastases developed by SW620 WT or HER3-KO cells. (J) Lactate secretion determined by ELISA. (K) Representative bioluminescent images of tumor-bearing mice and Kaplan–Meier overall survival of mice with orthotopic CRC liver metastases developed by SW620 cells. (L) IHC staining and quantification of PKF2 phosphorylation (S483) in human CRC and PDAC matched primary tumors (blue) and liver metastases (red). Each dot represents one biologically independent human sample. (M) Correlation between HER3 and PFK2 phosphorylation activations in human CRC and PDAC tumors, determined by *Spearman’s rank correlation coefficient.* Data in A—E, I, J, and L presented as mean ± SEM. \**p<0.01,* \*\**p<0.001,* \*\*\**p<0.0001,* \*\*\*\**p<0.00001,* by t-test. *p* value in K determined by log-rank test.

Seahorse metabolic flux analysis also showed lower OCR following HER3 activation **(Fig. 3C, Supp Fig. 3)**, consistent with increased glycolytic flux. HER3 activation further increased glucose consumption and lactate production, both of which were reversed by HER3 inhibition **(Fig. 3D, E)**.

Phosphofructokinase 2 (PFK2) is a key regulator of glycolysis. [16]. Immunoblotting showed that HER3 ligand stimulation increased PFK2 phosphorylation at serine 483 in CRC cells through distinct ligand-dependent kinase pathways. Specifically, recombinant human NRG1 (rhNRG1) activated the canonical AKT pathway, whereas recombinant human LRG1 (rhLRG1) signaled through RSK **(Fig. 3F).** Pharmacologic kinase inhibition revealed distinct ligand-specific signaling dependencies: rhLRG1-induced PFK2 S483 phosphorylation was completely abrogated by RSK inhibition (RSKi) while remaining insensitive to AKT blockade (AKTi), whereas rhNRG1-stimulated P-PFK2 S483 was suppressed by AKTi but unaffectedby RSKi **(Supp Fig. 4A).** These pathways converged on PFK2, providing a mechanism by which distinct HER3 ligands may enhance glycolytic flux in liver metastatic tumors.

**Figure 4.**
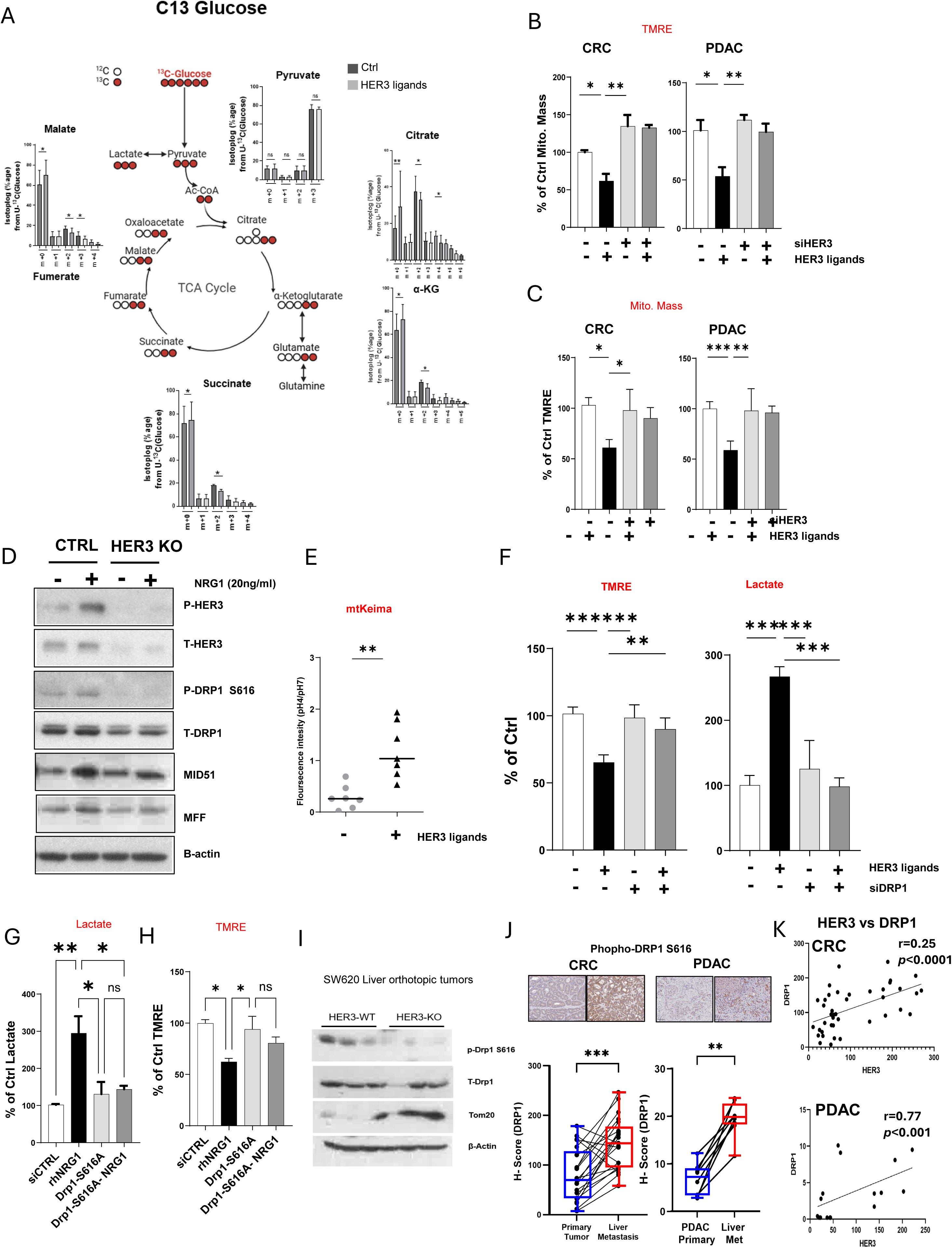
DRP1 signaling mediates HER3-induced metabolicdependencyfor mitochondrial OXPHOS. (A) Cancer cell glucose influx determined by ^13^C-glucose labeling and LC-MS analysis (B) Mitochondrial membrane potential determined by tetramethylrhodamine ethyl ester (TMRE) staining. (C) Mitochondrial mass determined by Mito Tracker staining. (D) Western blotting show levels of indicated proteins in SW620 human CRC cells with desired treatments. (E) Mitophagy determined by mtKeima reporter assays for mitochondrial pH shifting towards acidic. (F—H) Mitochondrial potential determined by TMRE, lactate secretion determined by ELISA. (I) Western blotting showing levels of indicated proteins in orthotopic CRC liver metastases developed by SW620 WT or HER3-KO cells. (J) IHC staining and quantification of DRP1 phosphorylation (S616) in human CRC and PDAC matched primary tumors (blue) and liver metastases (red). Each dot represents one biologically independent human sample. (K) Correlation between HER3 and DRP1 phosphorylation activations in human CRC and PDAC tumors, determined by *Spearman’s rank correlation coefficient.* Data in A—C, E—H, and J presented as mean ± SEM. \**p<0.01,* \*\**p<0.001,* \*\*\**p<0.0001,* \*\*\*\**p<0.00001,* by t-test.

Small interfering RNA (siRNA)-mediated HER3 knockdown prevented downstream PFK2 S483 phosphorylation (**Fig. 3G).** At the same time, pharmacological HER3 inhibition reduced PFK2 enzymatic activity **(Supp Fig. 4B).** To confirm that HER3 sustains PFK2 activation *in vivo*, we analyzed SW620 liver orthotopic tumors derived from HER3-WT and HER3-KO cells. Consistent with our *in vitro* signaling findings, HER3-KO liver tumors showed markedly diminished p-PFK2 S483 levels without changes in total PFK2 (T-PFK2) expression **(Fig. 3H)**. Correspondingly, enzymatic assays revealed an approximate 50% reduction in PFK catalytic activity in HER3-KO tumors compared with HER3-WT **(Fig. 3I)**, establishing that HER3 is required to maintain PFK2 phosphorylation and glycolytic enzyme activity within the hepatic metastatic niche.

To test the functional contribution of PFK2 S483 phosphorylation, we generated a phosphorylation-deficient S483A mutant [17]. The levels of of PFK2 S483A significantly decreased lactate production relative to wild-type PFK2 (**Fig. 3J**). In *a* CRC liver metastasis model, overexpression of PFK2 S483A (PFK2 Mut) significantly prolonged survival compared with PFK2 WT or control (**Fig. 3K**), supporting a role for this phosphorylation event in cancer progression. Consistent with these findings, IHC of paired CRC and PDAC primary tumors and liver metastases showed increased P-PFK2 S483 staining in liver metastases **(Fig. 3L).** Supporting our mechanistic findings that HER3 drives PFK2 phosphorylation and activity, linear regression profiling across primary vs liver metastases tumors determined consistent positive correlations between HER3 and PFK2 levels **(Fig. 3M)**, reinforcing the biological relevance of the HER3– PFK2 axis.

### HER3 activation promotes DRP1-dependent mitochondrial remodeling

The Warburg effect, characterized by preferential use of glycolysis over OXPHOS even when oxygen is available, is a hallmark of many cancers. However, the mechanisms linking oncogenic signaling to reduced mitochondrial activity remain incompletely understood [18]. To examine how HER3 couples increased glycolysis with decreased OXPHOS, we performed ^13^C-labeled glucose tracing. HER3 activation reduced labeling of tricarboxylic acid (TCA) cycle intermediates, suggesting decreased glucose carbon flux into mitochondrial metabolism **(Fig. 4A)**. In parallel, tetramethylrhodamine ethyl ester (TMRE) staining showed areduction in mitochondrial membrane potential following HER3 ligand stimulation in CRC and PDAC cells **(Fig. 4B).** Measurements of mitochondrial mass by Mito Tracker also showed reduced mitochondrial content in HER3-activated cells (**Fig. 4C, Supp Fig. 5**).

Drp1 is a GTPase that regulates mitochondrial dynamics, with phosphorylation at serine 616 linked to mitochondrial fission and cancer progression [19, 20]. To further assess HER3-dependent mitochondrial fission, we treated control (CTRL) and HER3-KO cells with or without NRG1. In CTRL cells, NRG1 increased Drp1 phosphorylation and the expression of MID51 and MFF; these responses were diminished in HER3-KO cells **(Fig. 4D**), which was further validated by blocking HER3 with inhibitors **(Supp. Fig. 6A**). Because mitochondrial fission can precede mitophagy, we next measured mitophagy flux using the mtKeima reporter assay. Indeed, activation of the HER3–Drp1 axis increased the mtKeima ratio towards low pH **(Fig. 4E)**, consistent with increased delivery of mitochondria to acidic lysosomal compartments. Consistent with a functional role for Drp1, siRNA-mediated Drp1 knockdown reduced HER3 ligand-induced lactate accumulation and reversed the decline in TMRE staining **(Fig. 4F**), together resulting in decreased cell growth **(Supp Fig. 6B)**,

To determine whether NRG1-driven metabolic reprogramming requires Drp1-S616 phosphorylation, we expressed a phospho-deficient Drp1 mutant (Drp1-S616A) prior to rhNRG1 stimulation[21]. While rhNRG1 markedly increased extracellular lactate and decreased mitochondrial membrane potential, Drp1-S616A expression completely abrogated both responses, maintaining lactate and mitochondrial membrane potential near baseline **(Fig. 4G, H)**. Immunoblotting of SW620 liver orthotopic tumors showed substantially lower p-Drp1 S616 levels, alongside increased Tom20 accumulation, in HER3-knockout tumors compared with HER3-WT controls **(Fig. 4I)**. Clinically, immunohistochemical H-scoring revealed significantly higher p-Drp1 S616 expression in matched liver metastases than in primary tumors for both CRC and PDAC cohorts **(Fig. 4J),** with HER3 and Drp1 levels positively correlating across both tumor types **(Fig. 4K)**. Together, these findings indicate that HER3 signaling promotes Drp1-S616 phosphorylation, coupling mitochondrial remodeling with enhanced glycolysis during hepatic metastasis. Together, these findings support a role for HER3 in promoting Drp1 S616-dependent mitochondrial fission and reduced OXPHOS in liver metastatic cancer cells.

## Discussion

Liver metastasis remains a major therapeutic challenge in advanced CRC and PDAC[22, 23]. In this study, we identify HER3 as a regulator of metabolic reprogramming in liver metastases, linking increased glycolysis and reduced OXPHOS to two downstream effectors, PFK2 and Drp1. This HER3-associated metabolic state creates a vulnerability to OXPHOS inhibition that can be exploited therapeutically. Combined HER3 and OXPHOS inhibition produced durable tumor-free survival in preclinical models of mCRC and mPDAC, providing a mechanistic rationale for further evaluating this therapeutic strategy.

One central finding is that HER3 supports glycolysis in the liver metastatic setting. The abundance of HER3 ligands, including NRG1 and LRG1 secreted by endothelial cells (ECs)[9], may selectively activate HER3 in disseminated tumor cells and promote metabolic adaptation to the hepatic niche. This interpretation is consistent with evidence that metastatic microenvironments impose distinct metabolic demands on cancer cells [24, 25] and that oncogenic receptor signaling connects extracellular cues to intracellular metabolic programs [26]. Our findings extend this model by suggesting that HER3 acts not only as a proliferative and pro-survival signal [8, 27] but also as a regulator of energy metabolism in liver metastatic cancer cells.

The HER3–RSK/AKT–PFK2 S483 axis provides a mechanistic basis for the glycolytic phenotype. HER3 activates downstream RSK and AKT signaling pathways, which converge on PFK2 to increase glycolytic flux and support metastatic progression. PFK2 is a bifunctional kinase/phosphatase that generates fructose-2,6-bisphosphate, a potent allosteric activator of phosphofructokinase 1 (PFK1), a rate-limiting glycolytic enzyme[16, 28]. Although RSK/AKT-mediated phosphorylation of PFK2 at S483 is known to enhance its kinase activity [17, 29], the involvement of HER3 in this regulatory process and in metastatic glycolysis has not been established. We show that PFK2 S483 phosphorylation occurs downstream of HER3 in vitro and is increased in CRC liver metastases. Moreover, the phosphorylation-deficient S483A mutant reduced lactate production and prolonged survival *in vivo,* supporting the functional importance of this modification. These findings identify the HER3–AKT/RSK–PFK2 pathway as a potentially targetable regulator of glycolysis in liver metastases.

In parallel with increasing glycolysis, HER3 signaling reduces mitochondrial function in association with Drp1 phosphorylation at S616. Drp1 is a key regulator of mitochondrial dynamics, and its activity is controlled by post-translational modifications. Phosphorylation at S616 promotes mitochondrial fission, which has been implicated in metabolic reprogramming and cancer progression[30] [31]. In our models, HER3 activation increased Drp11 S616 phosphorylation and reduced mitochondrial membrane potential and mass; HER3 inhibition or expression of Drp1 S616A reversed these effects. Elevated P-Drp1 S616 levels in patient liver metastases further support the relevance of this pathway. Thus, the HER3–PFK2 and HER3–Drp1 axes appear to act in parallel, increasing glycolytic flux while limiting mitochondrial respiration to reinforce the glycolytic phenotype of liver metastatic cells.

An important translational implication is that HER3 inhibition not only suppresses glycolysis but also exposes an increased dependence on OXPHOS in liver metastatic cancer cells. Combining HER3 inhibition with AG120 or metformin produced strong synergy *in vitro* and tumor-free survival in orthotopic mCRC and mPDAC models, with additional validation in PDX models. Activity in two cancers with distinct genetic backgrounds suggests that this strategy may extend to other HER3-expressing liver metastases. In addition, AG120 and metformin are clinically available drugs with established safety profiles, which may facilitate further evaluation of these combinations.

Several questions remain. AG120 and metformin have effects beyond direct mitochondrial targeting, and the specific OXPHOS complexes contributing to the observed synthetic lethality remain to be defined. Although our data implicated Drp1 S616 phosphorylation downstream of HER3, the kinase or kinases directly responsible for this modification within the HER3 signaling cascade also require clarification. Finally, the immunological effects of this combination therapy warrant investigation. Both glycolysis and mitochondrial metabolism influence the tumor immune microenvironment, and cancer cell metabolic reprogramming may alter immune cell function in the liver metastatic niche.

In summary, HER3 supports metastatic growth in the liver by promoting glycolysis and suppressing OXPHOS through pathways involving PFK2 S483 and Drp1 S616 phosphorylation, respectively. HER3 inhibition exposes an OXPHOS dependency that can be exploited through combination therapy, producing durable tumor control in preclinical mCRC and mPDAC models. These findings provide a mechanistic basis for HER3-mediated metabolic reprogramming in liver metastases and support further investigation of combined HER3 and OXPHOS inhibition in metastatic gastrointestinal cancers.

## Materials and Methods

### Cell Culture

The pancreatic ductal adenocarcinoma (PDAC) cell lines CFPAC-1, BxPC-3, and KPC were cultured in Dulbecco’s modified Eagle’s medium (DMEM) supplemented with 5% fetal bovine serum (FBS) and 1% penicillin–streptomycin. Cells were maintained at 37°C in a humidified atmosphere containing 5% CO2. HER3 knockout (KO) or knockdown cell lines were generated using CRISPR–Cas9-mediated gene editing (KPC) or shRNA (CFPAC-1), respectively, and validated by immunoblotting.

### Reagents

Sapitinib (MCE), a pan-HER inhibitor, was used as previously described (citation). AG120 (AstaTech), an IDH1 inhibitor, was used as previously described (citation). For *in vitro* experiments, metformin (MCE) was dissolved in sterile water at the indicated concentrations. IACS-010759 (MCE) and Lixumistat hydrochloride (MCE) were dissolved in DMSO at the indicated concentrations.

### Conditioned medium (CM)

Cancer cell-conditioned medium and liver sinusoidal endothelial cell (LSEC)-conditioned medium were prepared and used as previously described (citation: Liver endothelium promotes HER3-mediated cell survival in colorectal cancer with wild-type and mutant KRAS). For experiments involving AG120, conditioned medium was prepared using low-glucose (2.5 mM), low-magnesium (0.08 mM) medium (citation: IDH1 inhibition potentiates chemotherapy efficacy in pancreatic cancer).

### RT-qPCR

RNA was extracted from cultured cells according to the manufacturer’s instructions (Invitrogen). Complementary DNA (cDNA) was synthesized using the High-Capacity RNA-to-cDNA Kit (Applied Biosystems; Cat. No. 4387406). RT-qPCR was performed using PowerUp SYBR Green Master Mix (Applied Biosystems) on a Bio-Rad CFX96 system.

### Polar metabolite extraction for LC-MS metabolomics

For analysis of primary tumor and liver metastatic tissues, approximately 100 mg of snap-frozen tissue was transferred to a 2-mL microcentrifuge tube (Fisher Scientific) preloaded with zirconium oxide beads (1.4 and 2.8 mm; Precellys). Ice-cold 80% (v/v) HPLC-grade methanol (500 μL; precooled to −80°C) was added to each tube. Samples were homogenized using a Precellys homogenizer (5,000 rpm, one 30-s cycle) and centrifuged at 14,000 × g for 10 min at 4°C. The supernatant was transferred to a fresh tube on ice and dried using a SpeedVac vacuum concentrator without heat. Dried metabolite pellets were stored at −80°C until analysis. Protein concentrations were measured in a 100-μL aliquot of the extraction supernatant using a BCA assay (Pierce) for normalization.

Targeted polar metabolomic profiling was performed at the Mass Spectrometry Core Facility at Beth Israel Deaconess Medical Center (BIDMC), Harvard Medical School, as previously described[32]. For heatmap visualization, metabolite abundances were standardized as Z scores, calculated as (raw value − mean)/standard deviation (SD). This approach allows comparison of relative metabolite levels across features with different absolute abundance ranges.

### Extracellular flux analysis (Seahorse XF assay)

Mitochondrial respiration and glycolytic function were measured using a Seahorse XF HS Mini Analyzer (Agilent Technologies). Cells were seeded at 10,000 cells per well in Seahorse XFp Cell Culture Miniplates (#103025-100) and cultured in standard growth medium for 24 h, followed by treatment with the indicated reagents for 48 h. Sensor cartridges (#103022-100) werehydrated and calibrated according to the manufacturer’s instructions. On the day of analysis, the medium was replaced with Seahorse XF Assay Medium (pH 7.4) supplemented with 10 mM glucose, 1 mM pyruvate, and 2 mM glutamine. Cells were equilibrated for 1 h at 37°C in a CO2-freeincubator before measurement.

For the Glycolysis Stress Test (Seahorse XFp Glycolysis Stress Test Kit; #103017-100), extracellular acidification rate (ECAR) was measured at baseline and after sequential injections of glucose (10 mM), oligomycin (1 μM), and 2-deoxyglucose (2-DG; 50 mM) to assess basal glycolysis, glycolytic capacity, and nonglycolytic acidification, respectively.

For the Mitochondrial Stress Test (Seahorse XFp Cell Mito Stress Test Kit; #103010-100), oxygen consumption rate (OCR) was measured at baseline and after sequential injections of oligomycin (1.5 μM), carbonyl cyanide-4-(trifluoromethoxy)phenylhydrazone (FCCP; 1 μM), and rotenone/antimycin A (0.5 μM each) to determine basal respiration, maximal respiratory capacity, and nonmitochondrial oxygen consumption, respectively. ATP-linkedrespiration, proton leak, and spare respiratory capacity were calculated from the resulting OCR traces.

Seahorse measurements were normalized to cell number per well, determined using Quant-iT PicoGreen (Invitrogen) after each assay. Data were analyzed using Wave Desktop software (Agilent).

### Cell viability assay (crystal violet staining)

For cell viability assays, cells were seeded at low density in 96-well plates and treated with the indicated drug combinations for 7 days. At the endpoint, medium was aspirated and cells were washed once with 300 μL of PBS per well. Cells were fixed and stained with 100 μL of 0.5% crystal violet in 30% methanol per well for 2 h (CFPAC-1, BxPC-3, and KPC) or overnight (SW620 and HCT-116) at room temperature. Excess stain was removed by submerging the plate in running water and gently agitating it three times. Plates were inverted onto paper towels and air-dried overnight. To quantify staining, 100 μL of DMSO was added to each well and incubated for 1 h on a rocker at room temperature to solubilize the dye. A 50-μL aliquot of the solubilized dye was transferred to a 96-well plate, and absorbance at 600 nm (OD600) was measured using a microplate reader.

### Drug combination synergy analysis

To assess the effects of HER3 knockout (KO) in combination with OXPHOS inhibitors, wild-type (WT) parental or HER3 KO cells were seeded in 96-well plates and treated with AG120 (0, 0.005, 0.05, 0.5, 5, and 50 μM), metformin (0, 0.5, 5, 50, 500, and 5000 μM), IACS-010759 (MCE; 0, 0.0005, 0.005, 0.05, 0.5, and 5 μM), or Lixumistat hydrochloride (0, 0.005, 0.05, 0.5, 5, and 50 μM).

To assess drug combinations involving HER3 inhibition with sapitinib, cells were seeded in 96-well plates and treated with dose matrices of sapitinib (0, 1, 5, 10, and 20 μM) and either AG120 (0, 1, 5, 10, and 20 μM) or metformin (0, 0.5, 5, 50, 500, and 5000 μM).

After 7 days of treatment, cell viability was determined by crystal violet staining. Percentage inhibition relative to vehicle-treated controls was calculated for each single-agent concentration and combination. Drug interactions were analyzed using SynergyFinder (SynergyFinder.org) and the Loewe additivity model. Scores greater than 10 were classified as synergistic, scores between −10 and 10 as additive, and scores below −10 as antagonistic. Mean synergy scores are reported for each drug combination matrix.

### Human colon cancer organoid

Primary human colon cancer organoids were generated from deidentified discarded tumor tissue collected under institutional review board-approved protocols at Case Western Reserve University from patients undergoing resection for colon adenocarcinoma. Samples were isolated as previously described [33], and cancer organoids were cultured using established protocols[34, 35]. Following thawing, cancer organoids were passaged twice into 24-well plates and maintained in complete growth medium for 3 days to allow recovery. On day 4, organoids were transferred to nutrient-deficient medium for 48 h, followed by treatment for 24 or 48 h before downstream analysis.

### Organoid viability

To assess organoid growth and density, organoids were cultured in 24-well plates with/without treatment. Bright-field images of organoids embedded in Matrigel domes were acquired using a Leica Mateo FL microscope with a 2.5x objective. QuPath was used to quantify organoid area (μm²) and the number of organoids per Matrigel dome. Organoids with an area smaller than 50 μm² were excluded.

### Statistical analysis

Unless otherwise indicated, quantitative data are presented as mean ± standard deviation (SD) from at least three independent biological replicates. Two-group comparisons were performed using unpaired, two-tailed Student’s t tests. Three or more groups were compared using one-way or two-way analysis of variance (ANOVA), as appropriate. Survival was analyzed by Kaplan–Meier with p values determined by log-rank tests. Statistical analyses were performed using GraphPad Prism 10 (GraphPad Software). Statistical significance was defined as p < 0.05 (*p < 0.05, **p < 0.01, ***p < 0.001).

---

**Supplement Figure 1.**
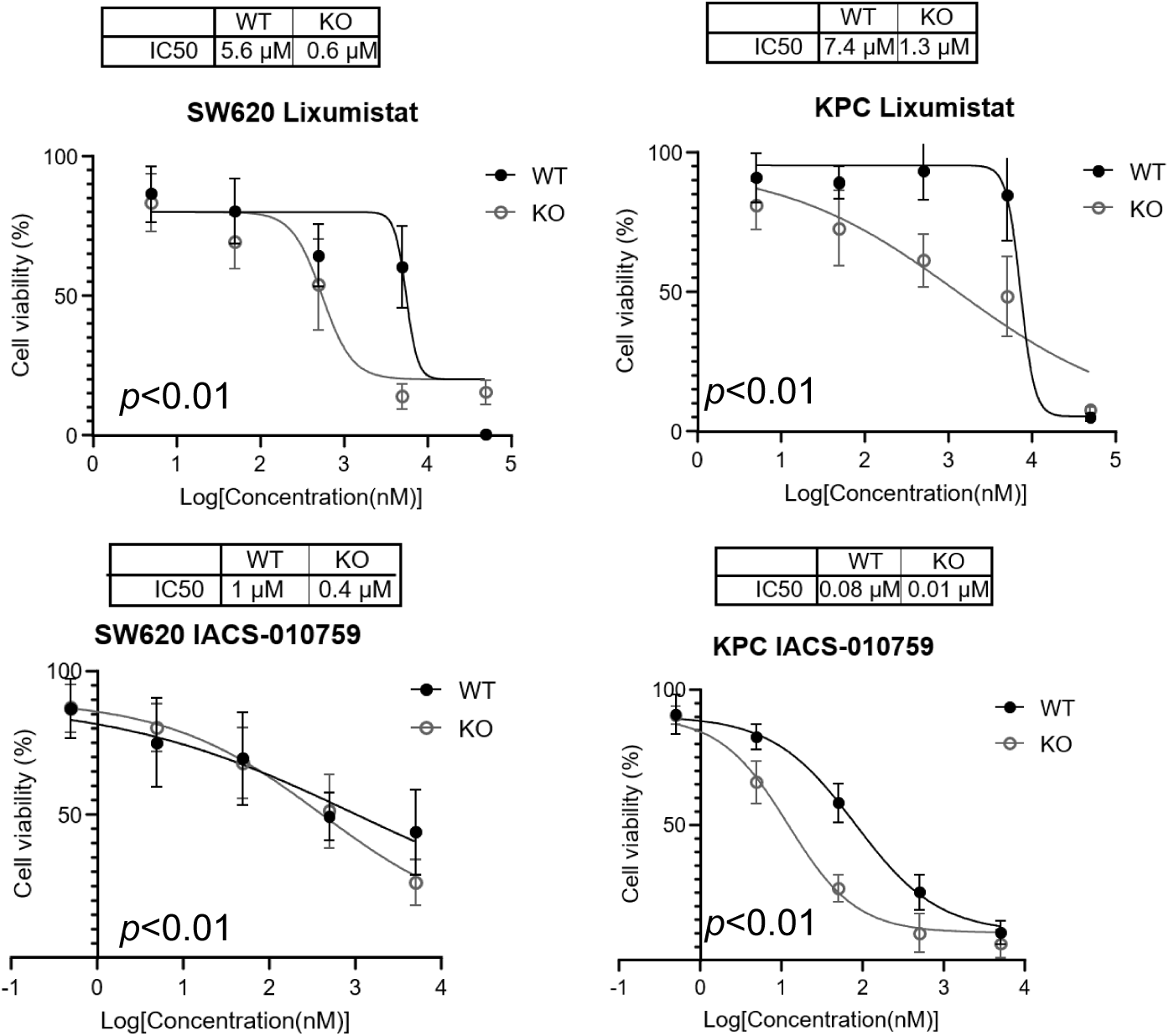
Combination of HER3 and OXPHOS Inhibitions. Relative growth and IC50 of WT and HER3 KO cells responses to metformin or IDH1 inhibitor AG120. *p* values determined by t-tests.

**Supplement Figure 2.**
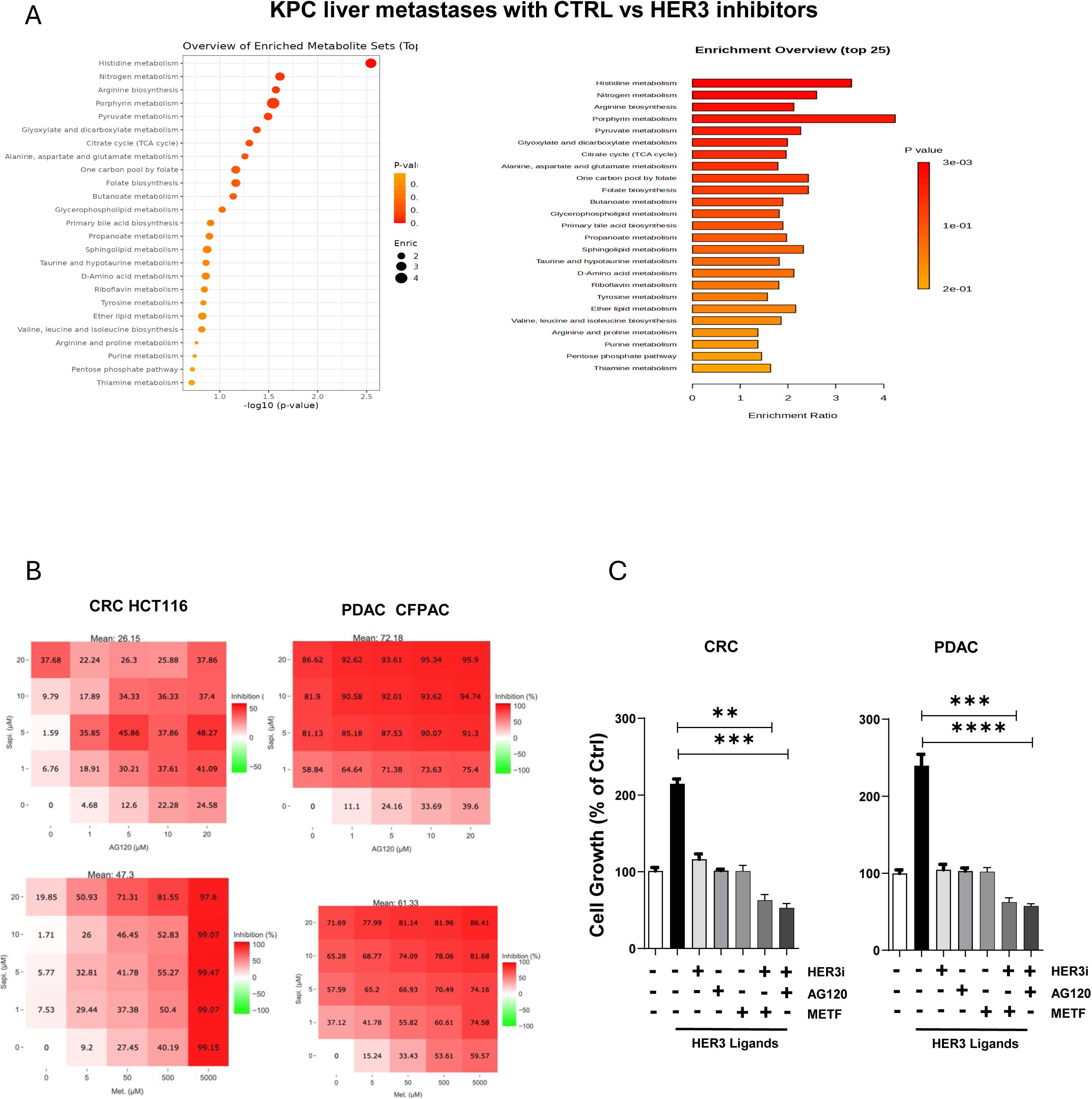
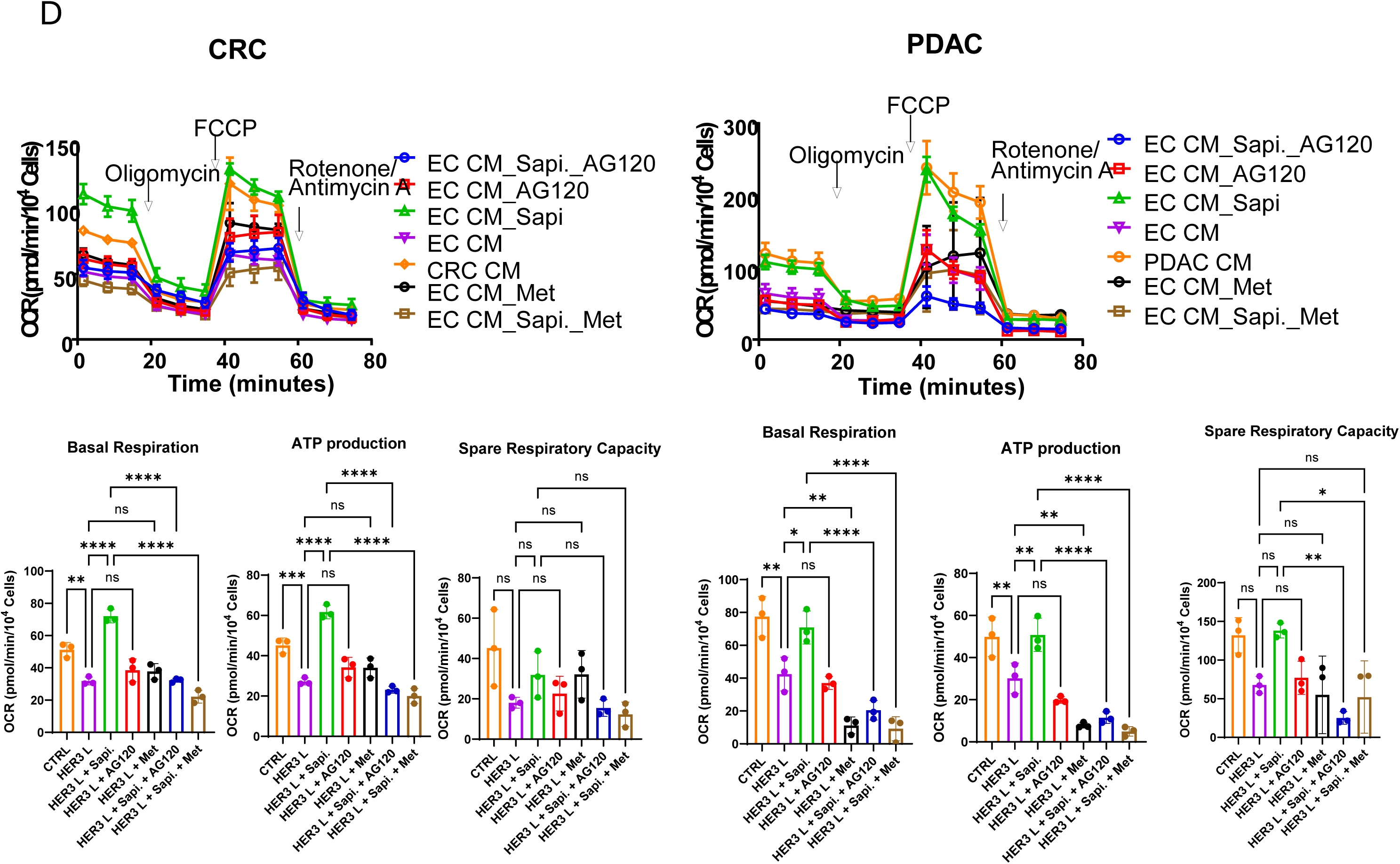
Combination of HER3 and OXPHOS inhibitions bydistinct inhibitors. (A) Heatmap and pathway analysis of metabolite changes between KPC orthotopic liver metastases treated without or with HER3 inhibitors sapitinib. (B) Cell survival from cells with the drug combinations in *in vitro* synergy assays. (C) Cell growth determined by colony formation. (D) Seahorse FX analysis of OCR with between-group comparisons for individual phases on the bottom. Data in C and D presented as mean ± SEM. \*\**p<0.001,* \*\*\**p<0.0001,* \*\*\*\**p<0.00001,* ns, not significant by t-test.

**Supplement Figure 3.**
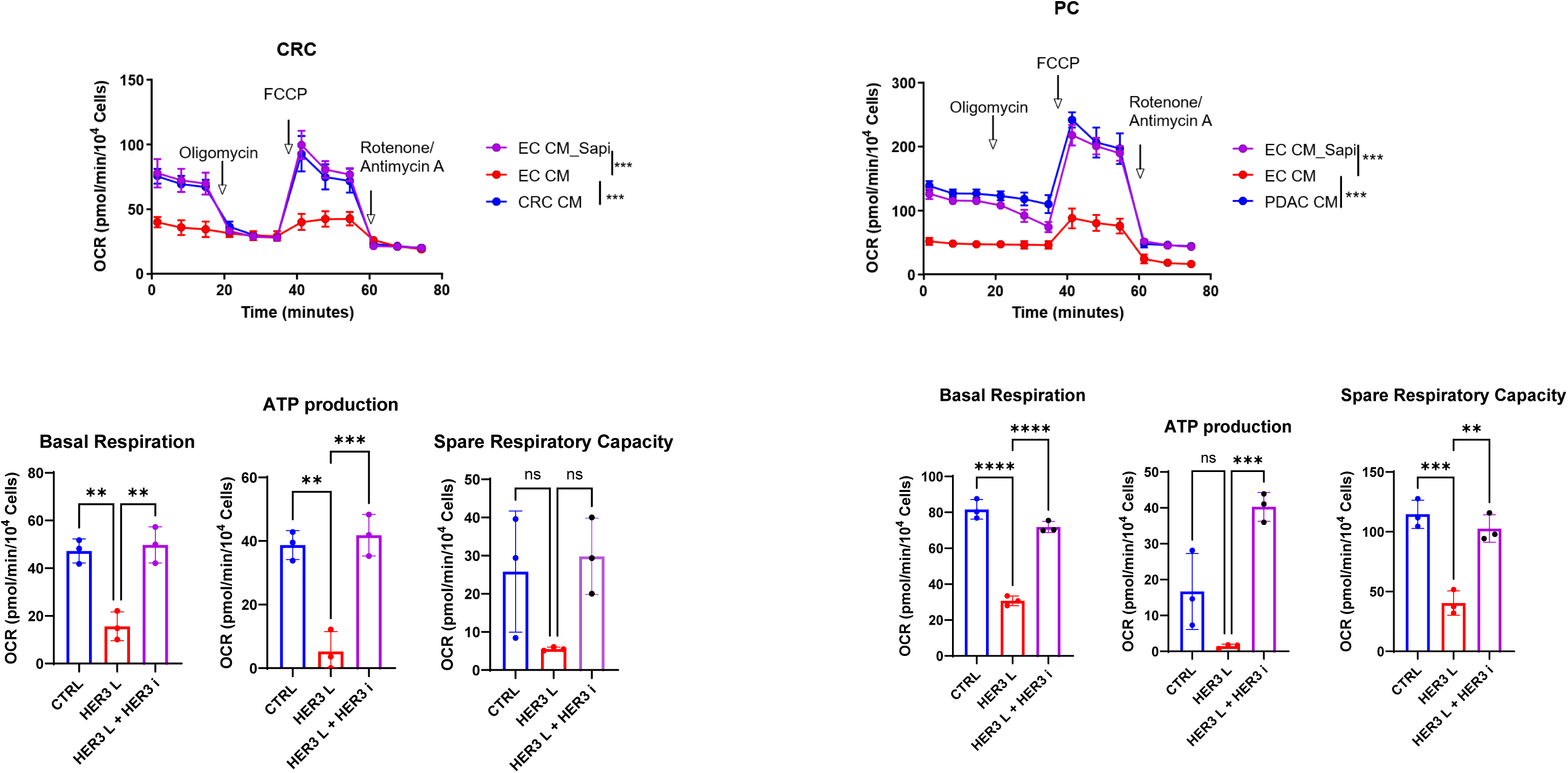
HER3 activation and inhibition affects cancer mitochondrial functions. Seahorse FX analysis of OCR with between-group comparisons for individual phases on the bottom. Data presented as mean ± SEM. \*\**p<0.001,* \*\*\**p<0.0001,* \*\*\*\**p<0.00001,* ns, not significant by t-test.

**Supplement Figure 4.**
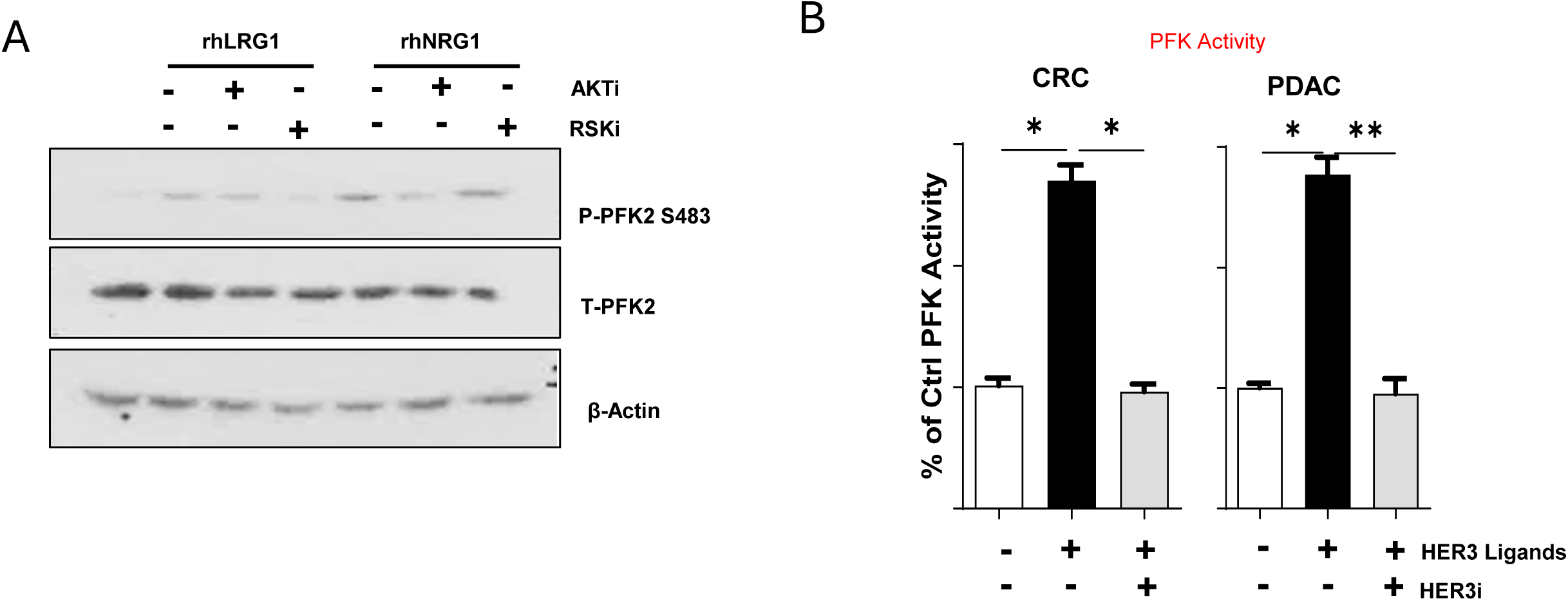
HER3 activation and RSK/AKT inhibition affect PFK2 activation. (A) Western blotting showing levels of indicated proteins in SW620 human CRC cells with drug treatments. (B) *In vitro* PFK enzyme activity determined by ELISA. Data presented as mean ± SEM. \**p<0.01,* \*\**p<0.001,* by t-test.

**Supplement Figure 5.**
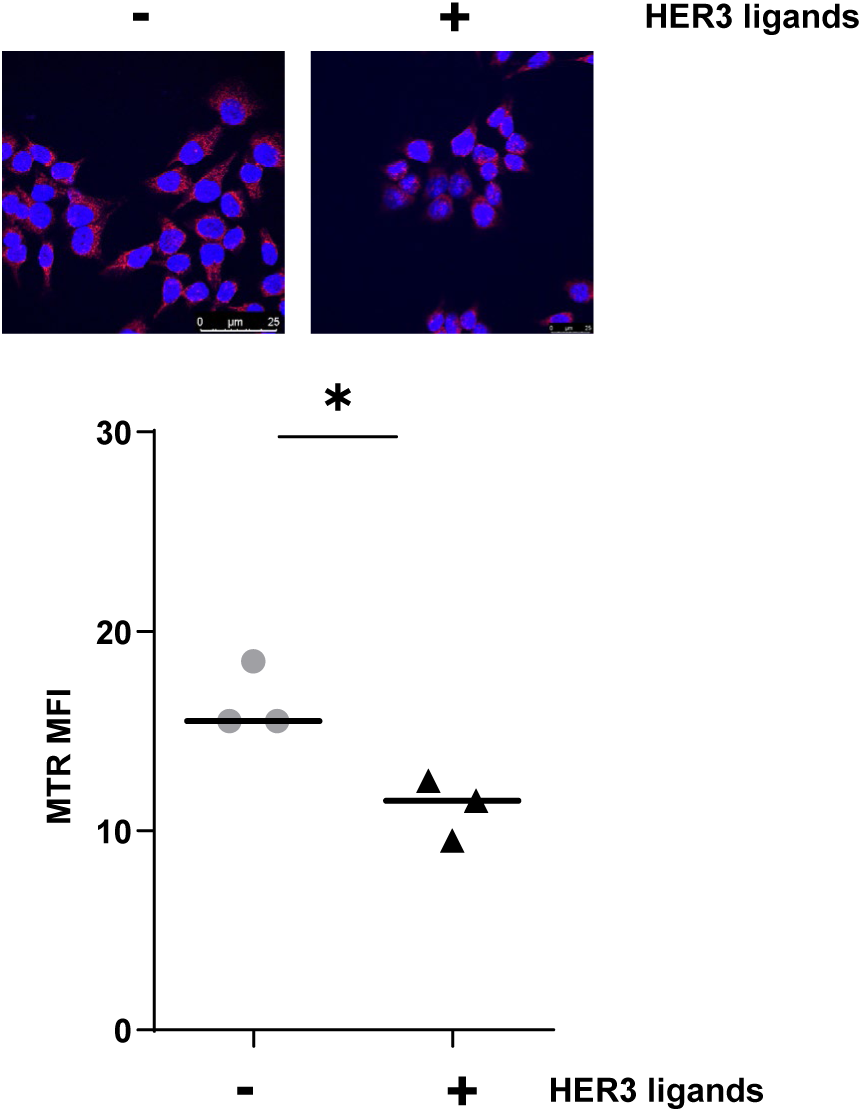
HER3 activation decreased mitochondrial mass. Representative images and quantifications of Mito Tracker staining (red) relative to DAPI nuclear staining (blue). Each dot represents mean of one independent experiment. Data presented as mean ± SEM. \**p<0.01,* by t-test.

**Supplement Figure 6.**
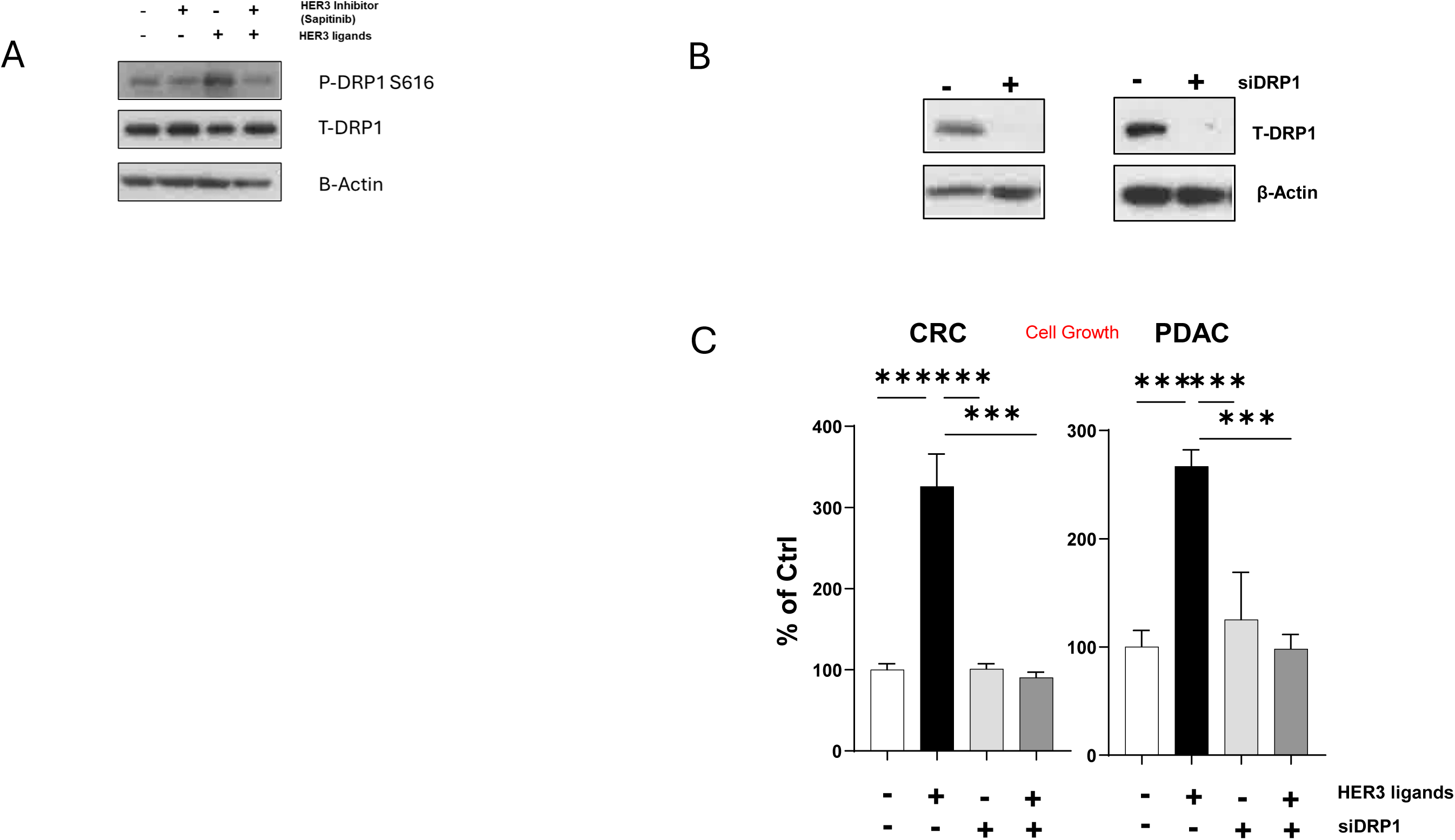
HER3 activation affect DRP1 activation. (A and B) Western blotting shows levels of indicated proteins in SW620 human CRC cells with desired treatments. (C) Cell growth determined by colony formation. Data presented as mean ± SEM. \*\**p<0.001,* \*\*\**p<0.0001,* \*\*\*\**p<0.00001,* by t-test.

